# Frequency-dependent organization of the brain’s functional network through delayed-interactions

**DOI:** 10.1101/754622

**Authors:** Abolfazl Ziaeemehr, Mina Zarei, Alireza Valizadeh, Claudio R. Mirasso

**Affiliations:** Department of Physics, Institute of Advanced Studies in Basic Sciences (IASBS), Iran; School of Cognitive Sciences, Institute of Research in Fundamental Sciences, Tehran, Iran; Institute of Interdisciplinary Physics and Complex Systems IFISC (CSIC-UIB), University Campus of the Balearic Islands, E-07122 Palma de Mallorca, Spain

## Abstract

The structure of the brain network shows modularity at multiple spatial scales. The effect of the modular structure on the brain dynamics has been the focus of several studies in recent years but many aspects remain to be explored. For example, it is not well-known how the delays in the transmission of signals between the neurons and the brain regions, interact with the modular structure to determine the brain dynamics. In this paper, we show an important impact of the delays on the collective dynamics of the brain network with modular structure; that is, the degree of the synchrony between different brain regions is dependent on the frequency. In particular, we show that increasing the frequency the network transits from a global synchrony state to an asynchronous state, through a transition region over which the local synchrony inside the modules is stronger than the global synchrony. When the delays are dependent on the distance between the nodes, the modular structure of different spatial scales appears in the correlation matrix over different specific frequency bands, so that, finer spatial modular structure reveal in higher frequency bands. The results are justified by a simple theoretical argument and elaborated by simulations on several simplified modular networks and the connectome with different spatial resolutions.

## Introduction

Brain is organized as a highly modular system at a hierarchy of spatial scales, from connectivity within a single cortical column to inter-areal brain-wide connectivity (Sporns and Betzel, 2016; Nicolini and Bifone, 2016; Zhou et al., 2006; Meunier et al., 2010; Park and Friston, 2013). Modules are sub-networks with high density of connections between the nodes within, and low density of connections between the modules (Girvan and Newman, 2002; Fortunato and Hric, 2016). This modular connectivity provides differential statistical inter-dependencies between the nodes inside and outside the modules and serves as a suitable substrate for functional specialization of the brain areas (Rubinov et al., 2009; Sporns, 2013; Lord et al., 2017). Presence of the long-range connections between a subset of nodes preserves the communication between the different modules, to promote functional integration (Tononi et al., 1994; Baum et al., 2017; Zamora-López et al., 2010).

Higher-order brain functions depend on the integration of local specialized processing in multiple spatial and temporal scales (Park and Friston, 2013; Bassett et al., 2011; Tononi et al., 1998; Cohen and D’Esposito, 2016). This integration takes place through a highly dynamic functional network: the intriguing property of the brain network is the ability to dynamically change the communication between the populations and the routes for information transfer despite to the fixed anatomy (Honey et al., 2007; Fries, 2005, 2015; Betzel et al., 2016). While time-averaged resting-state functional networks resemble the substrate structural network, there are considerable deviations from the structural networks when time-resolved, task-evoked functional networks are considered (Honey et al., 2007). It is hypothesized that the modular and hierarchical structure of the brain is particularly suited for providing such a diverse adaptive dynamics, but it is not well-known how the brain can spontaneously switch between the different functional patterns of communication on top of an almost static structural substrate (Park and Friston, 2013; Fries, 2005; Hutchison et al., 2013).

Interaction between the constituent nodes in the realistic distributed complex systems is not instantaneous. In brain circuits, in particular, the finite speed of signal transmission and synaptic processing time introduce a time delay in the interactions between neurons and between brain regions. According to experimental studies, the transmission delays in the mammalian brain has a wide range from milliseconds to tens of milliseconds (Ringo et al., 1994; Nakagawa et al., 2014; Ghosh et al., 2008; Asl et al., 2018). This indicates that the time delay can be comparable with some other important temporal scales in the nervous system, such as the membrane time constant, the period of gamma oscillations, and the effective window of time-dependent plasticity rules (Gray and Singer, 1989; Grenier et al., 2001; Abbott and Nelson, 2000); therefore not negligible. In the past few decades, the effects of time-delay interactions have received considerable attention. It has been shown that the presence of delay can substantially enrich the dynamics of physical and biological systems and lead to various phenomena such as multi-stability, clustering, frustration, and enhanced synchronization (Ermentrout and Kopell, 1998; Lee et al., 2009; Kim et al., 1997; Esfahani et al., 2016; Dhamala et al., 2004). In this study, we have shown how communication delays affect the dynamical interdependencies between the brain regions and give rise to a frequency-dependent functional network. We show that the pattern of the information transfer in neural circuits can be tailored by manipulating the frequency of the constituent nodes.

Transmission delays in the brain are heterogeneous and depend on the distance between connected regions (Swadlow and Waxman, 2012). Here we have explored the effect of heterogeneous delays on collective dynamics of the brain and phase relation between different brain regions, connected through the complex hierarchical-modular structure reported in the connectome datasets (Hagmann et al., 2008). To systematically explore the role of the various parameters and in particular to highlight the role of the distance-dependent delays, we began with a simplified synthetic modular network and systematically added the complexity to the network to end with the connectome.

We first provided a simple analytical approach (Wu and Dhamala, 2018; Woodman and Canavier, 2011; Esfahani and Valizadeh, 2014) to show that with delayed interaction, the connection between two oscillating components can be synchronizing or desynchronizing depending on the frequency of the oscillation. Through simulations on a synthetic modular network with homogeneous delays, we then showed that the local (intra-module) and the global (inter-module) coherence depend on the frequency of the nodes and there is a narrow transition frequency range over which the global order parameter is oscillatory and a stronger synchrony is observed within the modules due to the stronger intermodule connectivity. With heterogeneous distance-dependent delays, compatible with the theoretical prediction, we found a wide frequency range where the short- and long-range connections are synchronizing and desynchronizing, respectively, and strong local synchrony was observed while different modules evolved out of phase. In this region, the global order parameter was no longer time-dependent but took a very small value, and the functional network extracted from the dynamical correlation between the nodes closely resembled the modular structure of the underlying network. The same effect was observed in a modular structure with two levels of hierarchy where the delays were accordingly chosen from a trimodal distribution. In this case, increasing the frequency, the synchrony at successive hierarchical levels vanished at different frequencies, resulting in the distinct frequency ranges over each, the functional network resembled the modular structure at a different hierarchy level.

We finally run the simulations on the network extracted from the human connectome with different spatial resolutions (Hagmann et al., 2008), where the delays were assumed proportional to the distance between the nodes. The connectome data showed a modular structure with two levels of hierarchy, but the parameters such as the strength and the probability of the connections, number of nodes in the modules and the communication delays were more distributed than those in simpler synthetic networks. Moreover, the brain networks were distinguished from the conventional modular networks by the presence of networks hubs which were densely connected through inter-modular long-range connections. We showed that the matrix of the correlations in the model of the brain networks changes with the frequency, and the patterns of local and global synchrony change with frequency as is predicted by the theoretical background and simulations on simplified synthetic networks, but with less distinction between different states. We have explored and discussed the possible role of distributed parameters, the role of hubs and rich-club organizations in the connectome by performing extensive simulations. Our results highlight the role of communication delays, and in particular distance-dependent delays, in the generation of dynamic functional network and flexible information transfer pattern in brain networks.

## Methods

### Dynamical model

Our model consisted of N phase oscillators coupled through a modular network with time-delayed interactions described by a generalized Kuramoto model (Yeung and Strogatz, 1999):

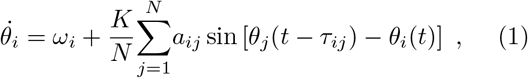

where, *θ*_*i*_ and *ω*_*i*_ *=* 2*πν*_*i*_ (*ν*_*i*_ being the frequency) are the phase and natural angular frequency of the *i*-th oscillator, respectively. *a*_*ij*_ are the elements of the adjacency matrix: *A*. *a*_*ij*_ = 1 if there is a directed link from the node *j* to *i* with a time delay *τ*_*ij*_; otherwise *a*_*ij*_ = 0. The parameter *K* sets the overall coupling strength.

### Data analysis

The degree of global phase coherence is quantified by using the instantaneous Kuramoto order parameter which is defined as 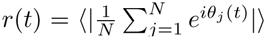, with 0 ⩽ *r*(*t*) ⩽ 1. *r*(*t*) = 1 (*r*(*t*) = 0) corresponds to the fully synchronized (incoherent) state (Kuramoto, 2003). Here 〈.〉 indicates averaging over different initial conditions. The time average of *r*(*t*) in the steady-state is represented by *R*. Moreover, to measure the degree of synchronization between any two nodes of the network, we use the correlation index defined as *σ*_*ij*_ = 〈cos[*θ*_*i*_(*t*) *− θ*_*j*_(*t*)]〉. Here *σ*_*ij*_ is an element of the correlation matrix *C* (Arenas et al., 2006). We determined *functional network* by the binarization of the correlation matrix. To this end, we introduced a threshold of *T* to convert the correlation matrix *C* to the functional network Σ.

The set of delayed differential equations (DDE) (Eq. 1) is solved numerically using adaptive Bogacki-Shampine scheme (Flunkert, 2011) with a minimum time step 0.05, absolute and relative error tolerance 10^*−*8^ and 10^*−*5^, respectively. The numerical integration was performed for the integration time *t*_*f*_ = 6500 for modular networks and connectome with 66 nodes and 10000 for hierarchical modular network and connectome with 998 nodes. In all the simulations we discarded *t*_*t*_ = 1000 as the transient time. The initial values of *θ*_*i*_ were randomly drawn from a uniform distribution in the interval [*−π, π*]. For simplicity, the natural frequencies of all the oscillators were considered identical.

### Network connectivity

To construct a modular network we considered a random network with *N* nodes which are grouped into *m* modules. In each module, nodes were connected with probability *P*_1_. The nodes in different modules were connected to each other with the probability *P*_2_ (*P*_1_ *> P*_2_).

To construct a hierarchical and modular network (HMN) we started with a modular network of *m*_0_ modules, each having *n*_0_ nodes. In each module, nodes were connected with probability *P*_1_ (first level of hierarchy). Nodes in different modules were connected with the probability *P*_2_ (*P*_2_ *< P*_1_) (second level of hierarchy). We add a copy of the above network and connect the nodes belonging to these two different sets with a probability *P*_3_, where *P*_3_ *< P*_2_ (third level of hierarchy). Generally speaking, to construct a network with *h* levels of hierarchy, we repeated the above procedure *h −* 1 times, with the hierarchical level-dependent probabilities, *pl* = *αq*^*l−*1^ (*l >* 1), where *α* is a constant, 0 *< q <* 1 and *q < P*_1_. The resulting network had 2^*h−*1^*m*_0_ modules and 2^*h−*1^*m*_0_*n*_0_ nodes (Moretti and Muñoz, 2013).

To construct the human connectivity matrix we used the human connectome data, down sampled from the high-resolution connection matrix (998 regions of interest (ROIs)) (Hagmann et al., 2008; Cabral et al., 2011). To downsample the matrix to 66 regions, all incoming fiber strengths to a target region were added and normalized by its region-dependent number of ROIs. The normalized coupling weights were assumed to be proportional to the number of tracts between the nodes and were controlled by an overall connection strength *K*. The graph constructed from the connectivity matrix had 574 edges, with an edge density of 27%, and contained 5 modules, its modularity was 0.45 and had an average degree of 18.

### Structure-dynamics similarity measure

To evaluate the degree of similarity between the binarized synchronization pattern matrix and the real modular/hierarchical network structure, we used normalized mutual information (NMI). It is a measure based on entropy that quantifies the amount of information shared by two random variables (here structural and dynamical patterns). The NMI was calculated by defining a confusion matrix **M**, for which the rows correspond to the structural modules and the columns to the dynamic modules found in a binarized correlation matrix. The elements of **M**, *M*_*ij*_, are the number of nodes in the real module *i* that appear in the emerging dynamic module

*j*. A measure of similarity between the partitions was then defined as (Danon et al., 2005)

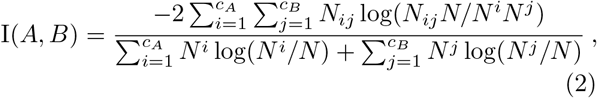

where the number of structural and dynamical modules was denoted by *c*_*A*_ and *c*_*B*_, respectively. The sum over row *i* and column *j* of the matrix *N*_*ij*_ was denoted by *N*^*i*^ and *N*^*j*^, respectively. The NMI gives results between 0 (no mutual information) and 1 (perfect similarity). In order to determine the dynamical modules in the binarized correlation matrix (functional network), we used the multilevel community detection algorithm proposed by Blondel et al (Blondel et al., 2008; Csardi and Nepusz, 2006).

## Results

We performed an extensive computational study on the pattern of the correlation matrix in a model of the human brain, where each region is characterized by a phase oscillator with a certain frequency. We focused on how the heterogeneous delays in the modular structure, affect the brain functional network at different frequencies. In the following, we first provide a theoretical background upon which the study is inspired. We then present the results of the simulations on the simplified multi-scale networks to highlight the role of delays and we then show how the results are applied to study the brain dynamics by operating the model on the human connectome networks.

### Theoretical background

We consider two phase oscillators with an identical frequency *ω*, coupled through a bi-directional Kuramoto-like sinusoidal delayed interaction function with strength *ϵ* and delay *τ*

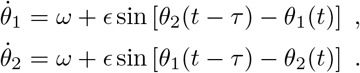

Assuming that the oscillators are weakly coupled, the evolution of the difference of the phases *φ* = *θ*_1_ − *θ*_2_ can be described by

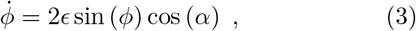

where *α* = *ωτ*. The phase locked solution determined by 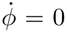 has two solutions 0 and *π*, where the former in-phase solution is stable if cos(*α*) *>* 0 and the latter anti-phase solution is stable if cos(*α*) *<* 0. So the nature of the connection depends on the normalized delay *ωτ*. Specifically, for *ωτ < π/*2 and *ωτ >* 3*π*/2 the connection is synchronizing, while for *π*/2 *< ωτ <* 3*π*/2 the connection is desynchronizing. In a network with fixed interaction delay, the nature of the connections can change due to the change in frequency. Of the interest in this study, is when the delays between the nodes are not equal, so changing the frequency, different connections change nature at different frequencies and at a given frequency some of the connections can be synchronizing, while the others have the reverse effect. In a modular network, it is reasonable to assume that the delays for long-distance inter-module connections are greater than those for local connections. This implies that increasing frequency from small values, first long-range connections change to desynchronizing and intermodule coherence is lost while local coherence within the modules persists. In the following, this hypothesis is examined and elaborated by extensive simulations on the synthetic simplified modular and hierarchical networks and in the realistic brain network.

### Modular networks with homogeneous and with distance-dependent delays

We first studied a network composed of phase oscillators divided into 3 modules. The connections between the modules were of the same strength, but less dense than the intra-module connections (Fig. 1a-b). Delays between any two nodes of the network were set equal, regardless of whether they are in a module or are in different modules. We then evaluated the overall coherence of the network using the global Kuramoto order parameter, for different values of coupling strength and mean intrinsic oscillator frequency (Fig. 1c). Consistent with the theoretical predictions, for small values of connection strength, increasing frequency the network switches from a global coherent state to the incoherent state through a transition state with partial synchrony, and with increasing the connection strength the incoherent region shrinks (Yeung and Strogatz, 1999). To clarify the dynamical properties of transition state we have shown the local (within the modules) order parameter along with the global order parameter for two different values of the connection strength in Fig. 1d-e. The plots show that in the transition region, which coincides in two spatial scales, the global order is slightly less than the local order.

**Figure 1:**
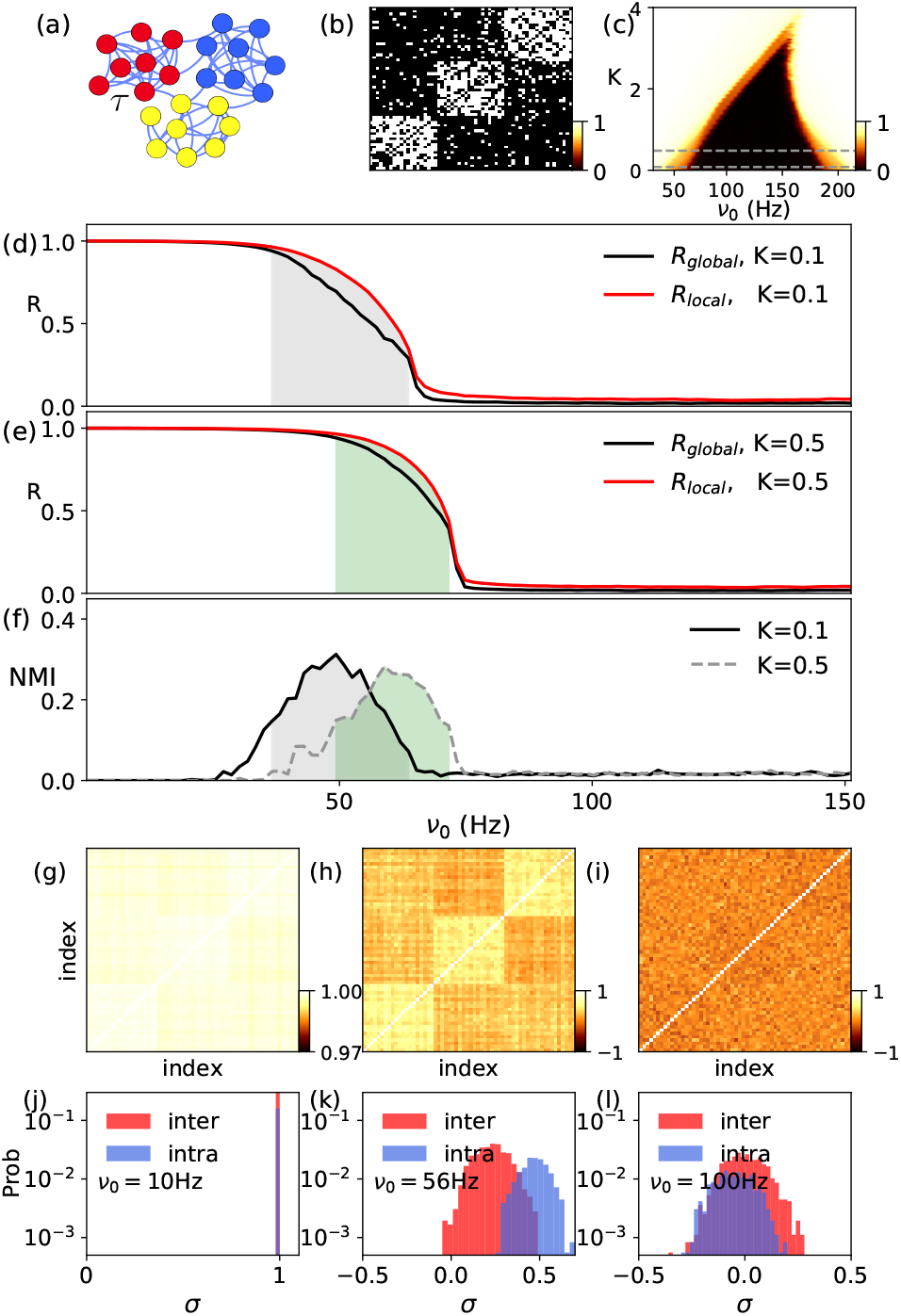
Modular network with homogeneous time delays. (**a**) The schematic illustration of a network with three modules. (**b**) The adjacency matrix of the modular network with *N* = 60, *m* = 3, (*P*_1_, *P*_2_) = (0.7, 0.1). (**c**) The stationary order parameter *R* for the Kuramoto model in the coupling strength *K*− average natural frequency *ν*_0_ phase space corresponding to the modular network with constant delay *τ* = 4. (**d, e**) The time average of global and local order parameters versus *ν*_0_ at couplings 0.1 and 0.5. The hatched area gives an estimate width of the boundary region. (**f**) The NMI versus *ν*_0_ for *K* = 0.1 and 0.5. (**g-i**) The correlation matrices (*C*’s) at the *K* = 0.1 and different *ν*_0_ = (10, 56, 100) Hz, corresponding to coherent, boundary and incoherent regions, respectively. (**j-l**) The distribution of correlation values corresponding to panel (g-i), respectively. The results were obtained by averaging over 50 different sets of network and initial phases.

This difference between the degree of synchrony in the local and global scales suggests that within the transition region, the modular structure could be inferred from the correlation matrix. To test this, we extracted correlation matrix for each value of frequency, of which, three typical ones are shown in Fig. 1g-i, representative of three regions of global synchrony, transition region, and asynchrony. Distribution of the correlation between the pairs within and between the modules for the same three frequency values are also shown in Fig. 1j-l. It can be seen that in the transition region, the correlation between the nodes inside and outside the modules take different mean values and the two distributions are slightly distinct. So in principle, it was possible to infer the modular structure of the network from the correlation matrix. To this end we extracted the functional network by binarizing the correlation matrix and evaluated the similarity between functional and structural networks, using the similarity measure NMI (see Methods). It can be seen in Fig. 1f that in the transition region, the two networks show a moderate level of similarity. The main constraint is that the threshold for the binarizing the correlation matrix should be chosen with care, for a suitable assignment of the functional link between the pairs inside and outside the modules. In the following, we will show that when the delays are distance-dependent, this constraint is relaxed and the modular structure has a more apparent effect on the functional network.

To explore the effect of distance-dependent delays, as the simplest exemplary network we considered a modular network with a bimodal distribution of delays, where the inter-module delays were greater than those for intra-modules (Fig. 2). The other parameters were the same as those of Fig. 1.

**Figure 2:**
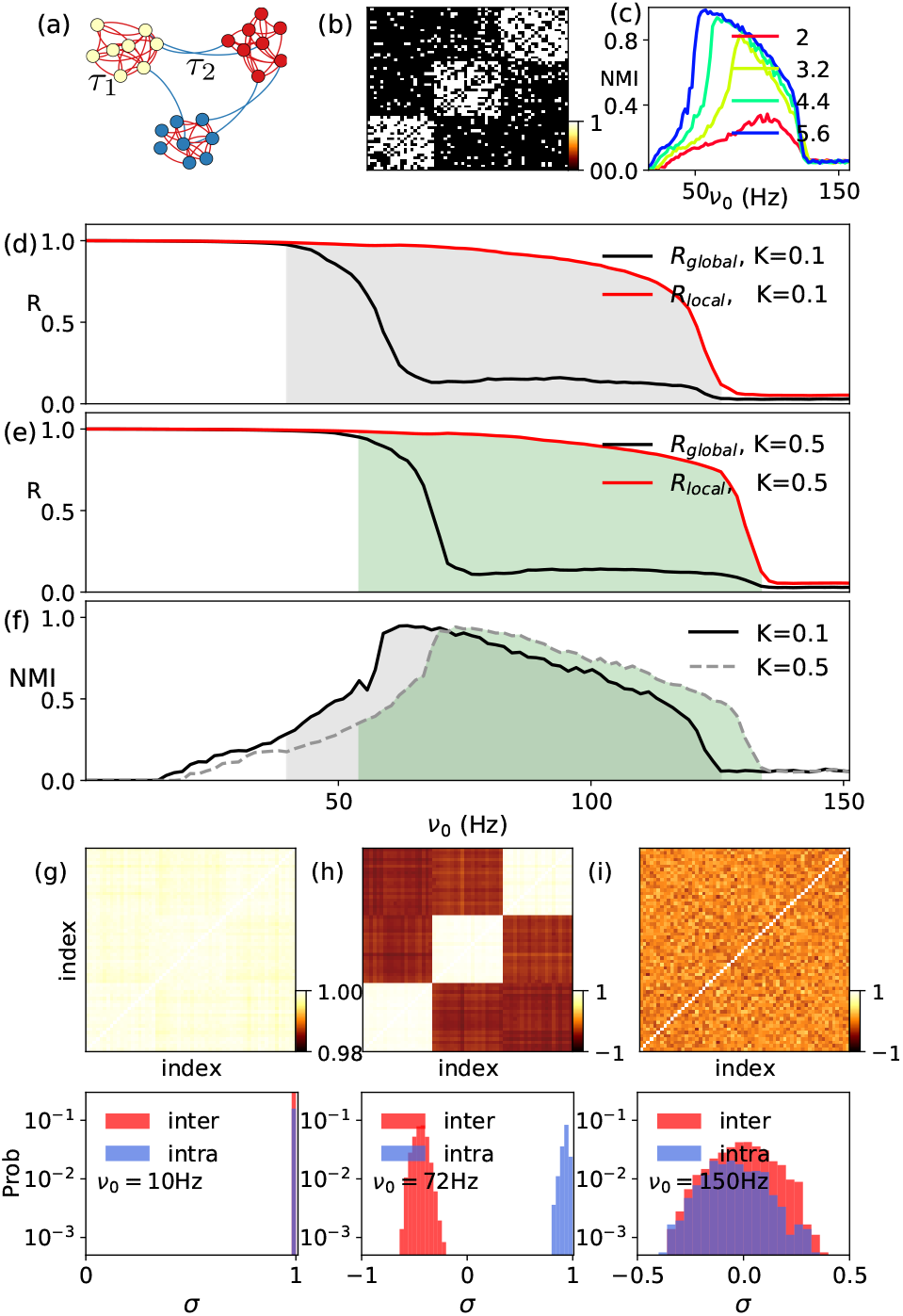
Modular network with distance-dependent time delays. **(a)** The schematic illustration of a network with three modules and bimodal interaction delay. **(b)** The adjacency matrix of the modular network with *N* = 60, *m* = 3, (*P*_1_, *P*_2_) = (0.7, 0.1). **(c)** The effect of different *τ*_2_ = 2, 3.2, 4.4, 5.6 and fixed *τ*_1_ = 2 on the NMI. **(d, e)** The time average of global and local order parameters versus *ν*_0_ at couplings 0.1 and 0.5, for connection matrix of (b) with(*τ*_1_, *τ*_2_) =(2, 4.3). The hatched area gives an estimate of the region of transition state. **(f)** The NMI versus *ν*_0_ for *K* = 0.1 and *K* = 0.5. **(g-i)** The correlation matrices (*C*’s) for *K* = 0.1 and *ν*_0_ = (10, 72, 150) Hz, corresponding to coherent, transition and incoherent regions, respectively. (**j-l**) The distribution of correlation values corresponding to panel (g-i), respectively. The results were obtained by averaging over 50 different realizations of network and initial phases.

The global and the local order parameters have been shown in Fig. 2d-e. We observe that increasing the frequency, over a wide region strong synchrony is observed within the modules, while global synchrony has taken a small value. We note that compared to the homogeneous delay case, here this region is much wider and the distinction between local and the global order parameters is quite bolder. The correlation matrices are shown in three panels of Fig. 2g-i and the corresponding distribution of the correlation of the pairs inside and outside modules Fig. 2j-l, indicate that compared to the homogeneous delay case, correlation across local and long-range connections are much more different. In particular, pairs in different modules show a negative correlation, indicative of anti-phase like dynamics.

The distinction between local and long-range correlation, in this case, suggests a closer correspondence between the structural and functional networks in the case of heterogeneous (distance-dependent) delays. To show this the binarized functional network was extracted from the correlation matrix and the similarity between the functional and structural connectivity was assessed using NMI similarity measure. We note that with heterogeneous delays, the results are independent of the choice of the threshold for extraction of the functional network from the correlation matrix, due to the well-separated distribution of the correlation, inside and between the modules. The results presented in Fig. 2f show that the range of frequencies over which the network structure and dynamics show similarity, gets wider and over this range the modular structure has a more clear reflection in the functional network, i.e., the NMI similarity index has a higher value compared to the previous ones with homogeneous delays (Fig. 2f). We also observe that the more differentiated the delays, the transition region widens and appears at lower natural frequency values (Fig. 2c).

To further clarify, and for a better comparison of the dynamics of the network with the homogeneous and heterogeneous delays, we have shown the time course of the phases and instantaneous Kuramoto order parameter for three different frequency values, corresponding to a coherent, incoherent, and the transition region for unimodal and bimodal delay distributions in Fig. 3. In the case of the unimodal delay distribution illustrated in Fig. 3a-c, in the transition region, a high (but not perfect) coherence is visible within the modules, while the three modules evolve out of phase. Different modules are not phase-locked and the phase difference between the modules is not fixed, leading to a time-dependent global order parameter Fig. 3b. Out of the transition region, the whole network is either synchronized Fig. 3a or desynchronized Fig. 3c.

**Figure 3:**
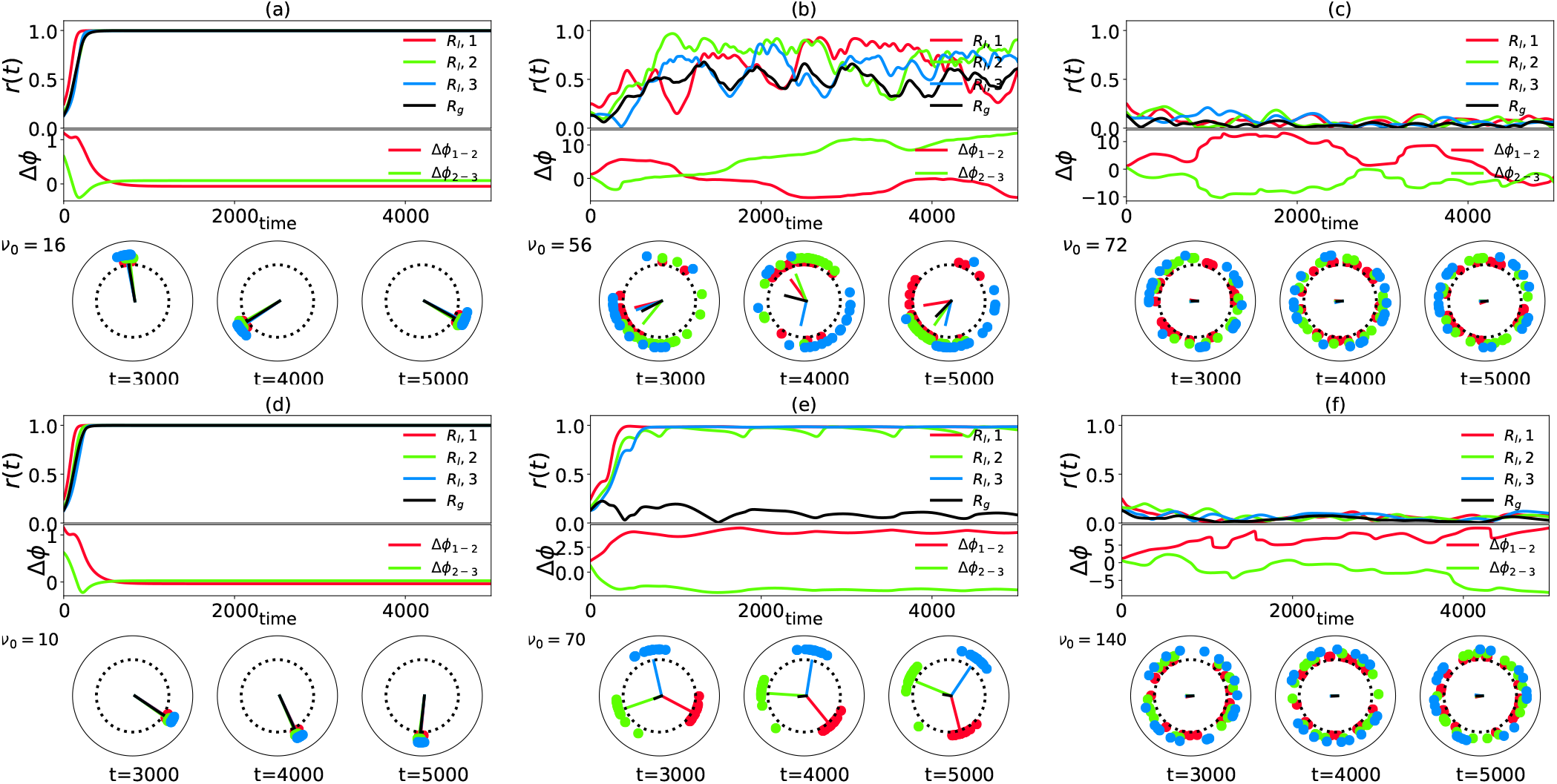
Dynamics of clusters for the modular networks. **(a-c)** and **(d-f)** show the dynamics of the clusters for a modular network with unimodal and bimodal delay distribution at different values of *ν*_0_ (the same network and delay as Fig. 1 and Fig. 2), respectively. **(Upper panel)** The global synchronization level (black) and the cluster synchrony levels (colored). **(Middle panel)** The time evolution of phase difference between clusters. **(Lower panel)** Three snap shots of node phases at given time. The colors are matched to the clusters. The corresponding order parameters for each cluster (colored) and the global order parameter (black) are represented inside the circles. Each set of **(a, b, c)** and **(d, e, f)** exhibit three phases of coherent, intermediate, and incoherent states, respectively.

The configuration of the phases in the three regimes with bimodal delay distribution illustrated in Fig. 3d-f shows that in this case, in the transition region strong synchrony is again found within the modules, while the three modules are phase-locked at the constant phase difference 2*π*/3. Therefore, while the synchronization in the modules is almost perfect, the fixed phase difference between the modules leads to a very low correlation between the modules and low global order parameter. These differences (compared to the unimodal delay distribution) lead to better separation of the correlation inside and between the modules in the case of distance-dependent delays.

As discussed in the theoretical background, the connection between a pair of oscillators with a given delay might be attractive or repulsive depending on the frequency, leading to in-phase or anti-phase locking, respectively (Ermentrout and Kopell, 1998; Deco et al., 2009; Sadeghi and Valizadeh, 2014). We distinguish two different mechanisms that can lead to the difference in synchrony level within and between the modules. In a modular network, because of the denser connectivity within the modules, the effective coupling, which is determined by the connection probability multiplied by the individual connection strength, is stronger inside the modules and the degree of the stability of the synchrony is different within and between modules. This means that the intra-module connection can lead to a stronger locking than the inter-module connections. In addition, different delays for intra and inter-module connections can result in a different (repulsive-attractive) nature of the individual links. Since the inter-module links have longer delays, they become repulsive for the lower frequency values, which makes it possible to lock the phases between the modules at the maximum possible phase difference, 2*π* divided by the number of modules (in the case of similar modules and the small number of modules). Although, the phase difference between the modules changes with the number of modules, a high local synchronization (with a small global order parameter) is obtained in this region. In summary, although high coherence and low coherence within and between modules are found in both uni- and bimodal delay distributions, different arguments may explain two cases. In homogeneous delay network, the lower connectivity between the modules leads to a lower correlation between the nodes in different modules in the transition region while for a bimodal delay distribution, the inter-module connections become desynchronizing in a range of frequency over which local connections are synchronizing. As a result, in the latter case, the level of correlation between the modules is much different than the correlation inside the modules, and the functional connectivity can more apparently reveal the underlying modular structure.

We also examined how the above results are consistent with respect to the changes in the different parameters of the network. This was an important question since the brain networks suffer from a substantial heterogeneity in the parameters. The results are presented in the supplementary figures (from S1 to S4). In general, the main result was valid, i.e., in all cases, the network transited from a globally synchronous state to an asynchronous state through a wide transition region with local synchrony, where the functional network was similar to the structural modular network. Still some effects were observed which are worth to be noted. In the figures S1 to S4 of the supplementary file we have shown how heterogeneity in the size of modules, presence of overlap between the modules, and heterogeneity in the delays inside and outside the modules can affect the dynamics of the modular networks. Interestingly, in all the cases a consistent phase relation between the modules no more existed in the transition region. This led to a time-dependent global order parameter and relatively smaller similarity between functional and structural networks. As we will see below all of these effects can be relevant in the study of the dynamics of the brain network.

### Hierarchical networks with distance-dependent delays

Brain networks show a modular structure in several hierarchical spatial scales (Zhou et al., 2006; Hagmann et al., 2008). As an advancement towards the real brain network, we applied the same analysis to the networks with more levels of structural hierarchy. Figure 4a-b shows the hierarchical network with three levels, constructed with (*m*_0_ = 1, *n*_0_ = 25, *P*_1_ = 0.9, *q* = 0.25 and *α* = 1). The dendrogram represents a hierarchical decomposition of the network which was generated by applying Ward’s agglomerative hierarchical cluster algorithm implemented in SciPy (Murtagh and Legendre, 2014; Oliphant, 2007). We considered only distance-dependent delays, chosen from a trimodal distribution where the delays increased with the connection level in the hierarchy. In this case, the globally coherent and fully incoherent states were separated in the parameter space by an intermediate region with local synchrony. Figure 4d shows the global order parameter and the local order parameters calculated over the modules in the two finer spatial scales. We observe that increasing frequency, first global synchrony is lost due to the decoherence of the two modules comprising the whole network, and then the synchrony within these modules is lost at larger frequencies since the modules in the finer spatial scale desynchronized. This was in accordance with our expectation that longer the connections (larger the delays), the synchrony is lost in smaller frequencies.

**Figure 4:**
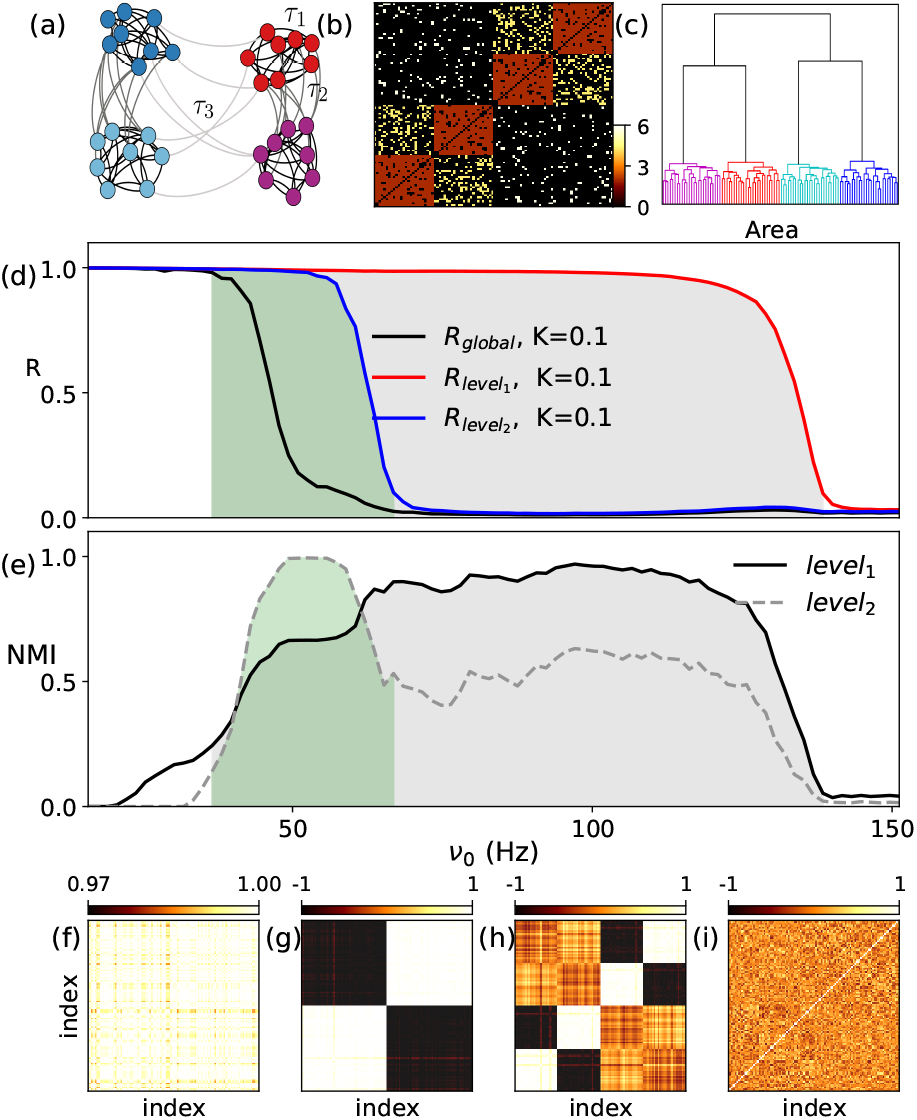
Hierarchical modular network with distance-dependent time delays. (**a**) A typical graph with hierarchical modular structure and heterogeneous coupling delays. (**b**) The adjacency matrix of the network. The colors show the values of the time delays for different levels of hierarchy ((*τ*_1_, *τ*_2_, *τ*_3_) = (2, 4.3, 5.7)). (**c**) The hierarchical clustering dendrogram corresponding to a network with *N* = 100 nodes and 2 levels of hierarchy with 2 and 4 clusters on each level. **(d)** The time average of global and local order parameters of clusters in level 1 and 2 versus *ν*_0_ at couplings 0.1 and 0.5, for connection matrix of (b). The hatched areas give an estimate of the boundary of intermediate state for each level. **(e)** The NMI measurement using the structural modules in the first and second levels for *K* = 0.1. **(f-i)** The correlation matrices (*C*’s) for *K* = 0.1 and different *ν*_0_ = (16, 48, 107, 145) Hz. The results were obtained by averaging over 50 different realizations of network and initial phases.

The NMI similarity measure illustrated in Fig. 4e shows the frequency ranges over which the functional network reveals the modular structure of the network in two levels. The second level (coarser scale) of the hierarchy is revealed over lower frequencies, while the first level (finer scale) became observable for higher frequencies. This point was more clearly apparent in the correlation matrices shown in the four panels of Fig 4f-i.

The phase dynamics of the nodes in the hierarchical network is illustrated in Fig. 5. For small values of frequency, all nodes oscillate synchronously. When increasing the frequency, first long-distance connections become repulsive because of longer delays. This cause the network to split into two subpopulations with antiphase dynamics. In each subpopulation, two modules belonged to a single cluster of the second hierarchy level (Fig. 5b). This is achieved in a frequency range where the second structural hierarchy level is revealed in the functional network. By further increasing the frequency, the intermediate-range connections also became repulsive and the four groups arrange their phases equidistantly at approximately the phase difference *π*/2 (Fig. 5c). This is accompanied by the range over which finer spatial scale of hierarchy appeared in the functional connectivity. It is notable that the phases of the two modules in a single cluster have a *π* phase difference because the repulsive connection between them was stronger than those in different clusters.

**Figure 5:**
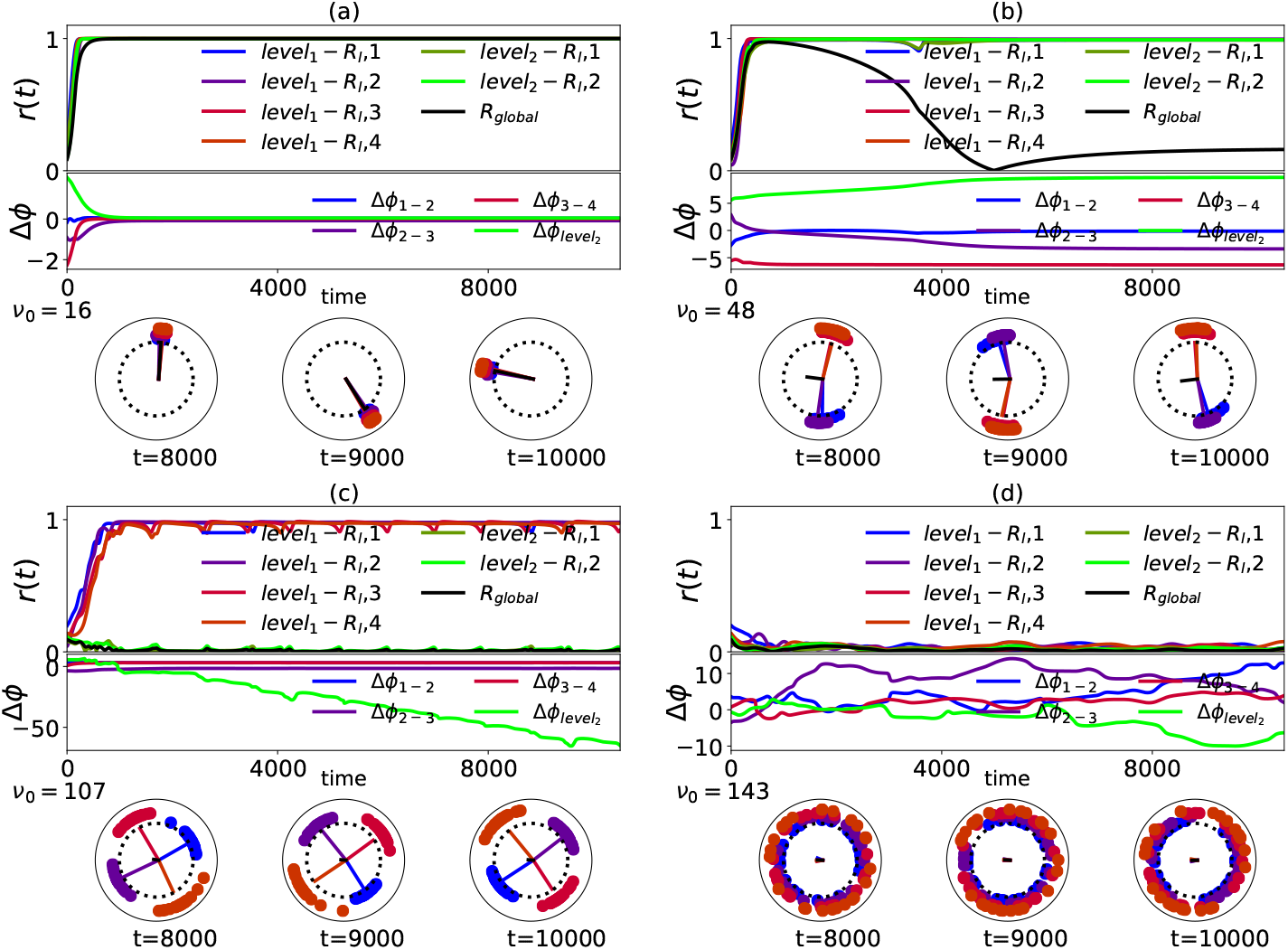
Dynamics of hierarchical networks with distance-dependent delays. The dynamics of the clusters for a hierarchical modular network at different frequencies is shown. **(Upper panel)** The global synchrony level (black) and the cluster synchrony levels (colored curves). **(Middle panel)** The time evolution of phase difference between clusters in level 1 and 2. **(Lower panel)** Three snapshots of node phases at the given times. The colors are matched to the clusters. The corresponding order parameters for each cluster (colored) and the global order parameter (black) are represented inside the circles. **a, b, c**, and **d** show four phases of globally coherent, cluster synchronized at second and first levels, and incoherent states, respectively.

#### Human connectome

In previous sections, it has been shown that the presence of distance-dependent delay of the interactions in a modular (and hierarchical) complex network, leads to a frequency-dependent pattern of synchrony in different spatial scales. Due to the finite velocity of signals along neurites and synaptic transmission times, interactions in the nervous system occur with delay (Jirsa, 2004; Nunez and Cutillo, 1995; Jirsa, 2009). In addition, the structural connections of the brain show a modular organization at different spatial levels. To check the validity of the mechanism presented above in the realistic brain networks, we run the model on the human connectome network. To construct the human connectivity matrix, we used human connectome data with 66 regions, downsampled from the high-resolution connection matrix (Hagmann et al., 2008) (see Methods). Fig. 6a shows the structural network, while Fig. 6b and c show the weighted connectivity matrix and the distance between the nodes, respectively. Similar experiments were also repeated on a higher resolution network with 998 nodes (Fig. S7).

**Figure 6:**
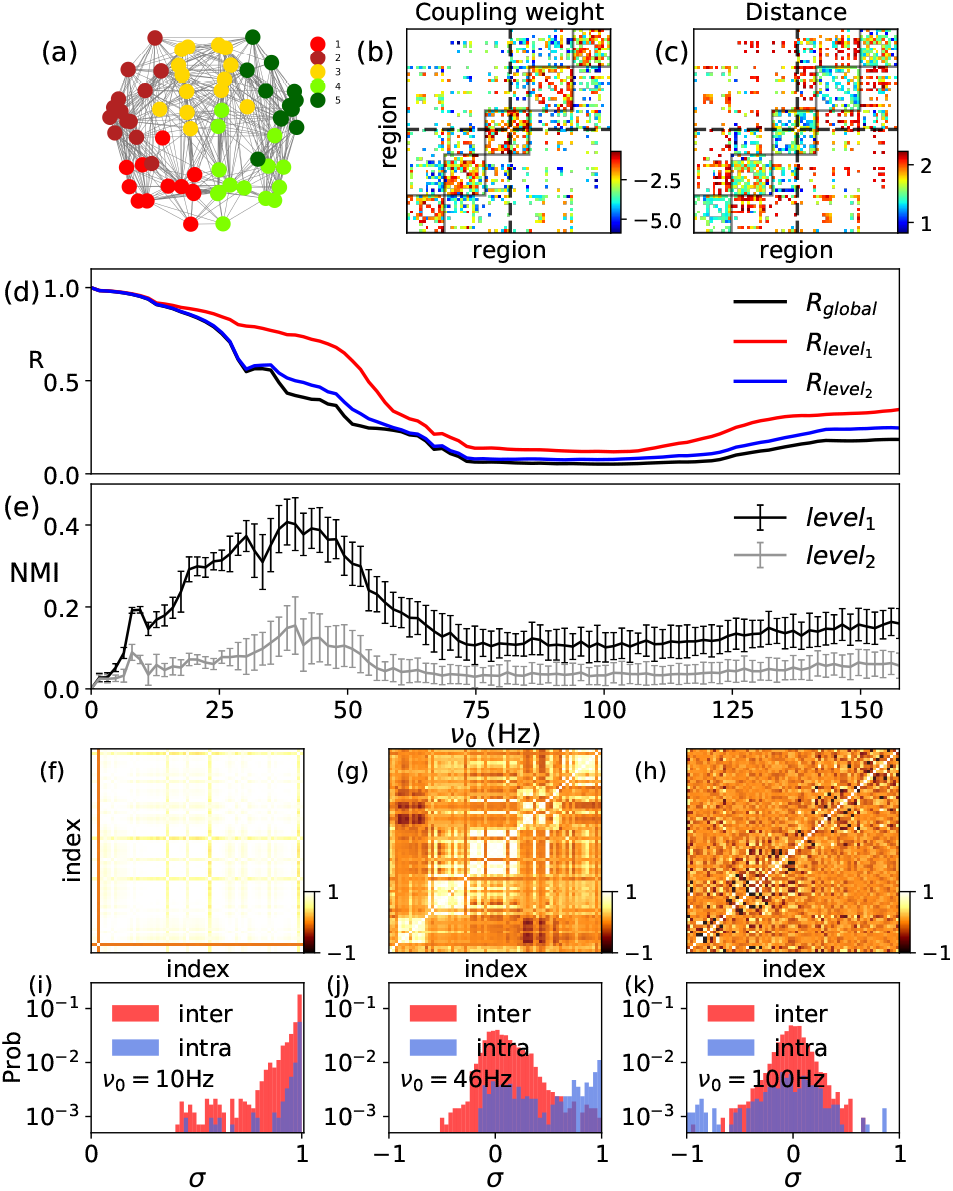
The human cortex. (**a**) The graph representation of the human cortex. The colors distinguish 5 communities. (**b**) The connection strength matrix, the color-bar has log_10_ scale. The weights were normalized so that 0 ≤ *W*_*ij*_ ≤ 1. (**c**) The distance between regions given as the average length of the fibers connecting pair of regions with log_10_ scale. **(d)** The time average of global and local order parameters of clusters in level 1 and 2 versus *ν*_0_ at couplings *K* = 30. **(e)** The NMI versus *ν*_0_ for *K* = 30. **(f-h)** The correlation matrices (*C*’s) at the same coupling *K* = 30 and different *ν*_0_ = (10, 46, 100) Hz. **(i-k)** The distribution of correlation values corresponding to panel (f-h), respectively. The results were obtained by averaging over 50 different sets of initial phases.

The parameters of the structural network are listed in the table 1. The connectivity matrix showed two levels of hierarchy where the whole brain was divided to two modules (corresponding to two hemispheres) in the coarser scale and to 5 modules in the finer scale. The main difference with the conventional hierarchical network studied above was that one of the modules in the finer scale is shared between two hemispheres (Fig. 6a). This network is reminiscent of the rich club organization with dense connectivity between a subset of nodes in different modules (Van Den Heuvel and Sporns, 2011; Harriger et al., 2012; Nigam et al., 2016; de Reus and van den Heuvel, 2013; Collin et al., 2013). Another important difference was that despite to our convention in the synthetic modular networks, the distance (and so the delay) between two nodes in different modules is not necessarily greater than those between the pairs inside the modules (see Fig. S5b,d). In other words, distribution of the delays for different spatial scales are overlapping, and inter-module connections include both short and long-range connections. This was more apparent in the second level of the hierarchy, i.e., the distribution of the delays connecting two nodes in single hemisphere almost spans the same range of delays compared with that of inter-hemisphere connections. Generally saying, all the parameter including the size of the modules, weight, and the delay of the connections were heterogeneous and were only partially distinctive for different spatial scales (Fig. S5 and table S1). This could affect the results as was studied in synthetic networks and was shown in the supplementary figures (Figs. S1-S4).

**Table 1:** Properties of each cluster in the human connectome with 66 nodes. Size shows the number of nodes, *e*_*in*_(*e*_*out*_) is the number of edges in (between) the clusters, *D* is the density of edges and *w*_*in*_(*w*_*out*_) is the normalized sum of edge weights in (between) the clusters. 〈*τ*_*in*_〉 and std(*τ*_*in*_) are the mean value of delay and standard deviation of delays, respectively.

| index | size | $e_{in}$ | $e_{out}$ | $D$ | $w_{in}$ | $w_{out}$ | $\langle \tau_{in} \rangle [\text{ms}]$ | $\text{std}(\tau_{in}) [\text{ms}]$ |
| --- | --- | --- | --- | --- | --- | --- | --- | --- |
| 1 | 12 | 34 | 116 | 0.52 | 0.17 | 0.03 | 6.45 | 4.3 |
| 2 | 13 | 57 | 123 | 0.73 | 0.16 | 0.02 | 9 | 6.6 |
| 3 | 14 | 61 | 159 | 0.67 | 0.19 | 0.04 | 6.4 | 4.5 |
| 4 | 14 | 48 | 158 | 0.53 | 0.17 | 0.03 | 6.6 | 4.6 |
| 5 | 13 | 47 | 98 | 0.71 | 0.16 | 0.02 | 8.2 | 6.1 |

To run the model on the connectome, we assumed that the communication delays are proportional to the length of the fibers connecting the nodes and assigned a constant average speed of 5m/s for signal propagation. Under these assumptions, the delay matrix took values between 0 and 40ms.

The global order parameter R and the local order parameters corresponding to the other two hierarchical scales were shown in Fig. 6d. Compatible with the previous results, when the delay increased, the system moved from coherence to the incoherent state through a relatively large transition region with partial synchrony. This was in analogy with the results on modular (Fig. 1 and 2) and hierarchical (Fig. 4) networks, however, the difference between the order parameters at different scales was not as bold as that in the synthetic networks, especially for the second level of the hierarchy. Based on the theory, that was because the delays are not completely differentiated for inside and outside the modules and for two hierarchical scales (see supplementary Fig. S5).

Functional network extracted from binarization of the correlation matrices at different frequencies was compared to the structural network using NMI measure, shown in (Fig. 6e). The similarity between the functional network and the finer scale modular structure showed a peak of around *ν*_0_ 40 Hz in the low gamma range is obtained (Pesaran et al., 2002; Henrie and Shapley, 2005; Bauer et al., 2007; Uhlhaas and Singer, 2006). But the second hierarchical state (corresponding to two hemispheres) showed a very low appearance in the functional network and almost at the same frequency which the first level appeared. It is worth to mention that in hierarchical networks we expected that the finer scale structure to reveal over higher frequencies, given the delays in two levels have distinct distributions. Here the distribution of the delays was not distinctive for the second level and accordingly, both the spatial levels appeared over the same range of frequencies. The exemplar correlation matrices presented in Fig. 6f indicate the appearance of the finer scale modular structure in the correlation matrix over the appropriate range of frequency. Similar results were obtained for high-resolution human connectome with 998 nodes, shown in supplementary Fig. S7.

Figure 7 contains more detailed information on the dynamics of the nodes in different regimes. For low frequencies, high values of global and local order parameters, indicate coherence within and between modules. For large frequencies, coherence is lost at the local and global levels. In the intermediate region, local-order parameters have larger mean values and less temporal variation than the global order parameter. The important difference here, compared to the simulated modular network (Fig. 3), was that the modules are not phase-locked to each other. This was the consequence of the wide distribution of the different parameters of the human connectome, where among them the distribution of the delay is the most decisive parameter that can destabilize perfect locking of the modules (see tables 1, S1). This was in agreement with our results presented in the supplementary material showing that heterogeneity destroys phase locking between the modules (Figs S1-S4).

**Figure 7:**
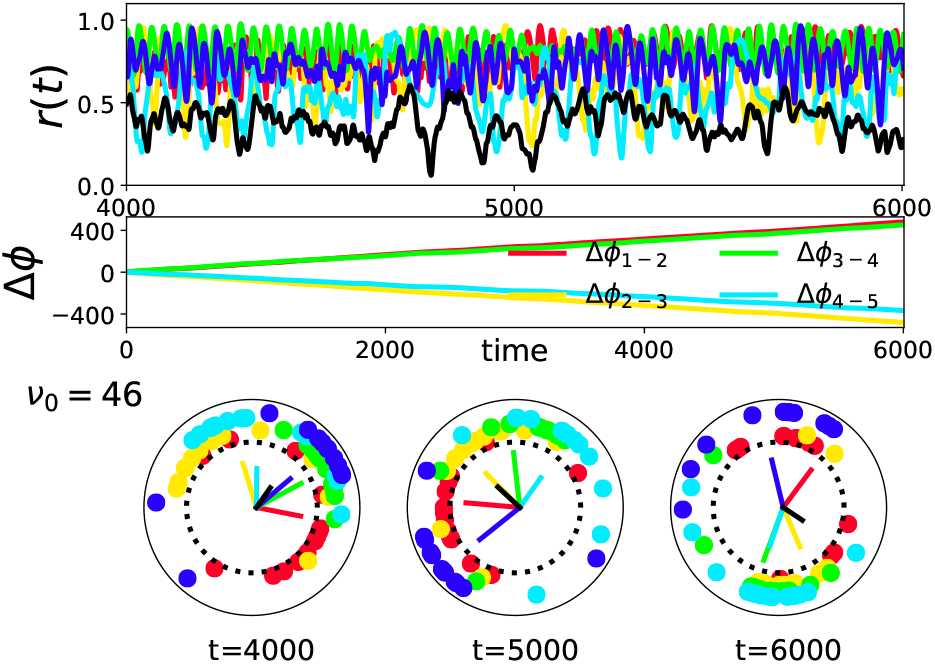
The cluster dynamics for the human connectome with 66 nodes at *ν*_0_ = 46 Hz. **(Upper panel)** The level of global synchrony (black) and the synchrony levels of the clusters (colored curves). **(Middle panel)** The time evolution of phase difference between clusters **(Lower panel)** Three snap shots of node phases at given time. The colors are matched to the clusters. The corresponding order parameters for each cluster (colored) and the global order parameter (black) are represented inside the circles.

## Conclusions

Impact of the structure of the neuronal networks on their dynamics has been extensively studied in recent years (Sporns et al., 2000; Honey et al., 2007; Hermundstad et al., 2013), but many aspects are not fully understood. The purpose of the present study was to investigate the role of time-delayed interactions and the modular structure in shaping the collective dynamics of the brain networks. In particular, we were interested in the role of distance-dependent delays on the correlation between oscillatory activities of different brain regions. To this end, we first simulated simple modular networks of phase oscillators and explored the role of delayed interactions. The main result was that the pattern of the synchrony was dependent on the frequency of the nodes. Increasing the frequency of the constituent oscillating nodes, the networks switched from the coherent to the incoherent state via a partially coherent state in which a stronger coherence was seen between the nodes within the modules compared to those in different modules. In the case of homogeneous synchrony, the transition region was narrow and the local and global coherence had a small difference. When the delays were distance-dependent, the transition region considerably widened and the difference between the coherence inside and outside the modules became quite significant: Almost perfect synchrony was maintained inside the modules while different modules dynamics was un- or anti-correlated. Over the suitable range of frequency, i.e., in this transition region, the modular structure of the network was reflected in the *functional network* deduced by the binarization of the networks correlation matrix. The similarity of the structural and functional network was almost perfect over a wide frequency range if the delays were picked from a bimodal distribution with two distinct peaks for inter- and intra-module connections.

We also run the model on top of the human brain connectivity matrix. It is known that the brain has a modular multi-scale structure and the neural signals do take a delay to be transmitted between any two nodes of the neural network, due to the finite speed of the signals (Aston-Jones et al., 1985; Antic and Zecevic, 1995). We observed that the modular structure can be revealed by the correlation matrix in a range of frequencies, but only a low (for coarser spatial scale) to moderate (for finer spatial scale) level of similarity was observed between the structural and functional networks, consistent with the previous results (Cabral et al., 2014). We attributed these results to the several differences between our synthetic modular networks and the brain network. One of the most important differences was the presence of a subset of nodes in different modules which are strongly connected together. This rich-club population(Harriger et al., 2012; Nigam et al., 2016; de Reus and van den Heuvel, 2013; Collin et al., 2013) could maintain correlation between the modules for a wider range of parameters. The next difference was that the distribution of the delays in the connectome was not quite distinctive for the links inside and outside the modules and there was considerable overlap between two sets of links. Moreover, heterogeneity in different parameters of the brain network such as the size of the modules, strength of the connections and connection probability could affect the results. Our extensive simulations on the variations of the synthetic modular networks showed that these considerations can be indeed an explanation for the observed results on connectome and their differences with the results on synthetic modular networks.

Our results are also noteworthy for their applicability to network inference in realistic networks. The first is that our method, like many other methods, is based on the transparency of the structure near the verge of coherent state or in the transition region over which partial synchrony can be observed in the collective dynamics (Villegas et al., 2014; Kaiser et al., 2007). Most of the other methods for the network inference are based on the varying an overall interaction strength to set the network in the transition region (Cabral et al., 2012). Although the strength of the synaptic connections in the brain changes due to the synaptic plasticity (Song et al., 2000; Tsodyks et al., 1998), these changes are too slow to explain the changes in the pattern of functional interactions which can take place in a few milliseconds (Biswal et al., 1995). Here we have shown that this could be achieved by varying the frequency band, given the connections bear non-zero delay. Notably, in case that the delay of the connections was specific for different hierarchical levels, the modular structure of the network became very apparently transparent in a wide range of frequency, and there was no need for a precise adjustment of the parameters to infer the structure from the functional network.

In the conventional models of synchronization, without time delays, the synchronization properties are independent of the mean frequency of the nodes and are simply determined by the connection strength and the disparity of the natural frequencies (Kuramoto, 2012; Acebrón et al., 2005). The presence of a delay introduces a new time scale into the equations of the system and makes them dependent on frequency (Yeung and Strogatz, 1999; Esfahani et al., 2016). Indeed, the interaction can force the nodes to an in-phase or an anti-phase state, depending on the delay and oscillation frequency (Wu and Dhamala, 2018; Li and Zhou, 2011). When the links have different delays, the identification of different links in the network can be different at a given frequency and it is possible to engage and disengage different nodes in the network in coherent communities with adjusting the baseline frequency. In this way the present study proposes that the heterogeneous delays play a decisive role in the state-dependent pattern of the coherence in the brain (Hutchison et al., 2013) and the flexible pattern of effective communication between the brain regions (Friston, 2011; Pariz et al., 2018).

