## Supplementary figures for "Frequency-dependent organization of the brain’s functional network through delayed-interactions"

### Supplementary information

#### 1 The Impact of various structural heterogeneities on the dynamics

As we can see in the **table 1**, **S1** and **figure S6**, the cortical modules have different sizes, densities and overlap with each other. Therefore, we study the effects of having modules with various sizes and densities or having overlapped-modules on the Kuramoto dynamics. Initially, we start considering networks with homogeneous modules. **Figure S1** determines which of the dynamical properties of a modular network change by changing the size of modules. In the transition region, we can find parameters that, at the stationary states, modules are almost phase-locked in equidistant angles, and the global order parameter is close to zero.

We also looked at overlapping modules. **Figure S2** shows the results obtained by considering overlapped modules. We can see that increasing the number of overlapping nodes leads to the larger amplitude oscillations in the order parameter of the modules and higher synchronization of the network. In fact, by increasing the number of overlapping nodes, anti-phase synchronization of the modules is converted to the in-phase synchronization. Therefore the dynamics can not reveal the structure of a widely overlapped modular network.

We continue our analysis by studying networks with heterogeneous modules. **Figure S1** treats modules of different sizes in a similar analysis. The results show that size heterogeneity destroys phase-locked states, which can be observed by growth and oscillation of the global order parameter in the transition region. It is interesting that broadening in size distribution of modules increases the amplitude of this oscillation. Also, this heterogeneity leads to a wider frequency region for module detection.

**Figure S3** shows that mixing modules with heterogeneous densities leads to similar results, i.e. large amplitude oscillations in the global order parameter for all parameters and a wider frequency region for module detection.

In summary, in the networks with homogeneous modules and time-delayed interactions, regardless of the size of modules, we can find a parameter in the transition region that the modules are almost phase-locked in equidistant angles, and global order parameter is close to zero. Interestingly, increasing the number of overlapping nodes converts this anti-phase synchronization to in-phase synchronization. On the other hand, increasing heterogeneous artifacts in the network structure, by mixing modules with different sizes or densities, destroys phase-locked states and makes order parameter growth and oscillations. Moreover, heterogeneity cause that the range of frequencies over which the network structure becomes transparent gets wider.

We also studied the heterogeneity in the time delay distributions. **Figure S4** shows that when we have homogeneity in the delay structure of the modules, whether we choose delays from uniform distributions or constant values, there are parameters that the modules are phase-locked and the global order parameter is zero. We can also see that heterogeneity in the delay distribution causes order parameter growth and oscillations for all parameters in the transition region. Therefore, in the real networks like connectome that we have heterogeneous modules, we will have order parameter oscillations and rhythms for all the parameters in the frequency region between coherent and incoherent states.

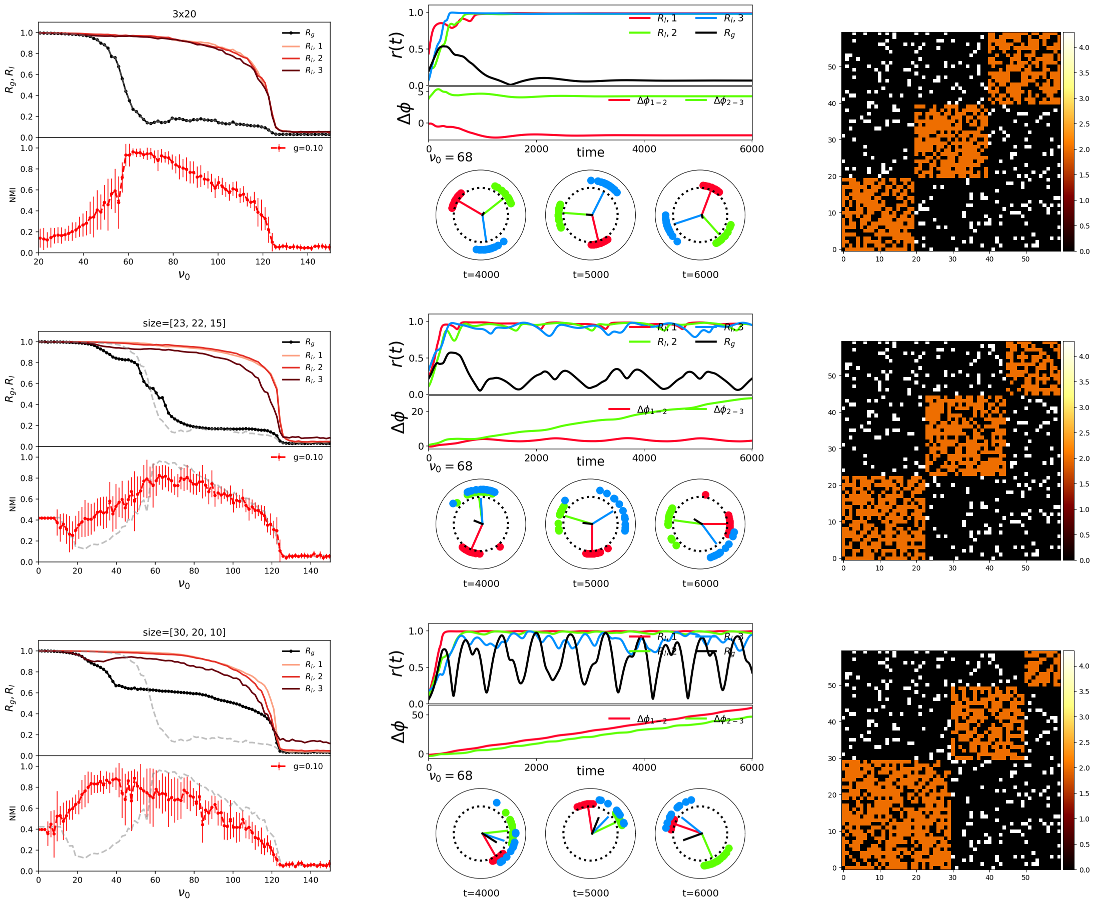


Figure S1: **The impact of heterogeneous module size distribution.** The results are for networks with N = 60 nodes and 3 modules contained (**Upper**) (20, 20, 20) nodes (**Middle**) (23, 22, 15) nodes and (**Lower**)(30, 20, 10) nodes. (**Left column**) represent the order parameters for the whole network and each module and NMI versus frequency. The results are averaged over 10 realizations. The gray dashed lines are the global order parameter and NMI for the upper row network and are plotted for the eye guide. (**Middle column**) show global and modules' order parameters and the phase differences between modules versus time. Three snapshots of the phase of the oscillators in the unit circle have been also shown. (**Right column**) indicate the connectivity matrices and color codes used to identify the values of time delays. Here, $(\tau_{in}, \tau_{out}) = (2, 4.3)$ and $(p_{in}, p_{out}) = (0.7, 0.1)$.

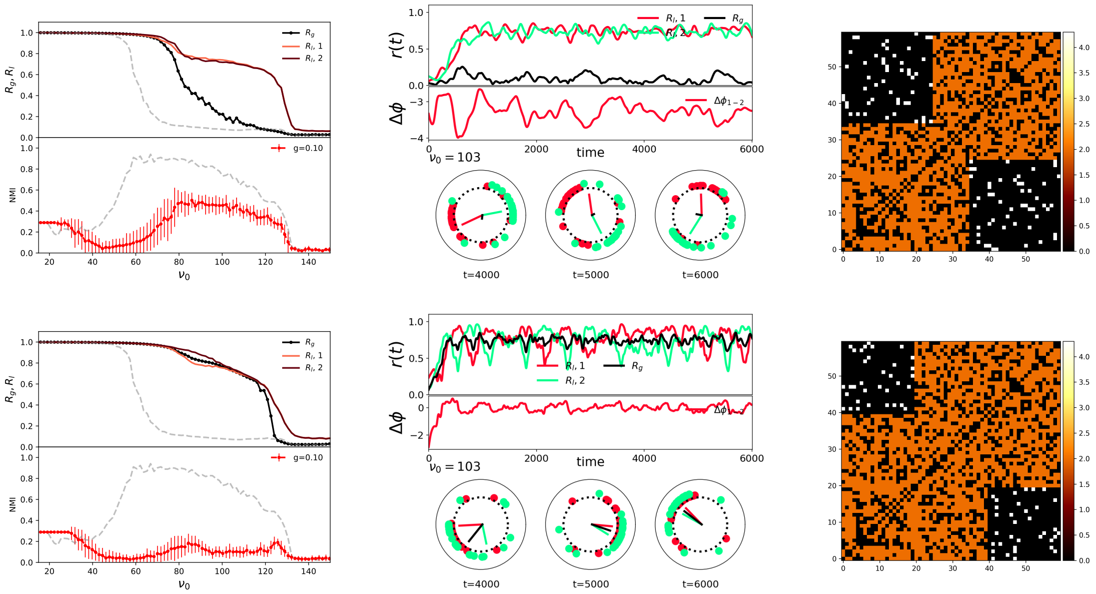


Figure S2: **The impact of overlapping nodes.** In the upper (lower) panels the networks have 60 nodes, 2 equal-sized clusters with 5(10) overlapping nodes. (**Left column**) Represent the order parameters for the whole network and each module and NMI versus frequency. The results are averaged over 20 realizations. The gray dashed lines are the global order parameter and NMI for the network with 2 non-overlapping equal-sized modules and are plotted for the eye guide. (**Middle column**) show global and modules' order parameters and the phase differences between modules versus time. Three snapshots of the phase of the oscillators in the unit circle have been also shown. (**Right column**) indicate the connectivity matrices and color codes used to identify the values of time delays. Here, $(\tau_{in}, \tau_{out}) = (2, 4.3)$ and $(p_{in}, p_{out}) = (0.7, 0.1)$.

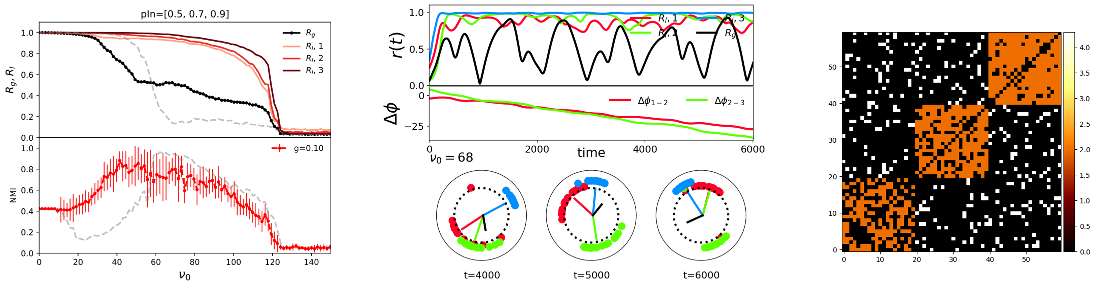


Figure S3: **The impact of heterogeneity in the edge density of modules.** (**Left panel**) shows the order parameter for the whole network and each module, and NMI versus frequency for networks with N=60 nodes and a different number of modules. The results are averaged over 10 realizations. The gray dashed lines are the global order parameter and NMI for the network with 3 equal-sized modules and $(p_{in}, p_{out}) = (0.7, 0.1)$ (the first row of figure S1) which are plotted for the eye guide. (**Middle panel**) shows global and modules' order parameters and the phase differences between modules versus time. Three snapshots of the phase of oscillators in the unit circle have been also shown. (**Right panel**) indicates the connectivity matrix and color codes used to identify the values of time delays. Here, $(\tau_{in}, \tau_{out}) = (2, 4.3)$, and the edge probabilities inside of modules are $(p_1, p_2, p_3) = (0.5, 0.7, 0.9)$ and $p_{out} = 0.1$.

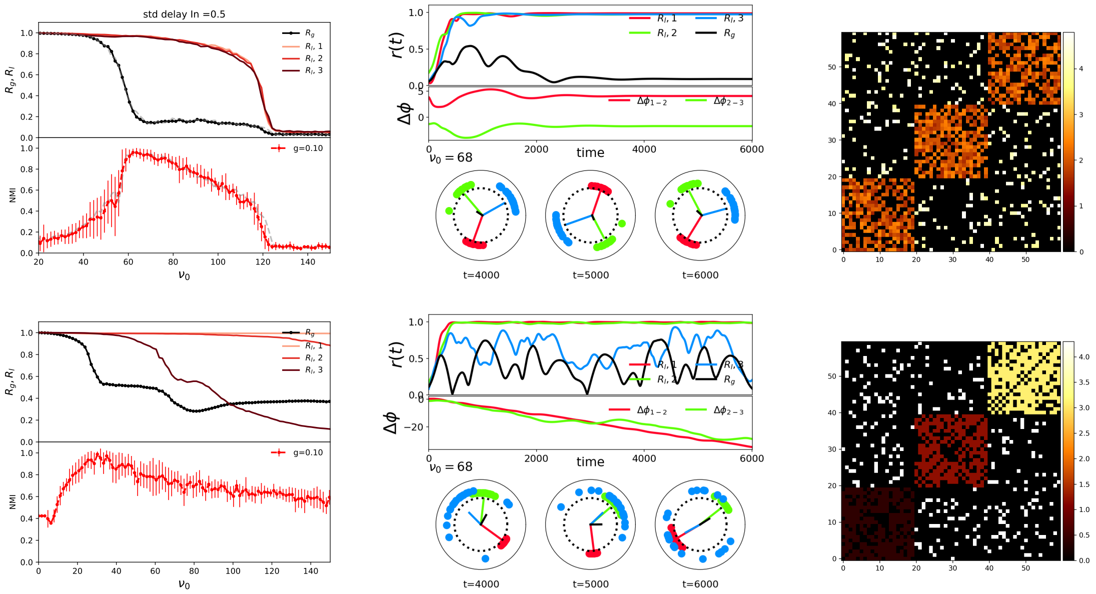


Figure S4: **The impact of heterogeneity in the distribution of delays**. (**Upper row**) shows the results when the time delays for the links inside and between the modules have been taken from the uniform distributions ($\tau_{in} \in [1.5, 2.5]$ $\tau_{out} \in [3.8, 4.8]$). (**Lower row**) represents the results when there is heterogeneity in the values of the delays for the links inside of different modules ($\tau_{in} = (0.4, 1.2, 3.1)$ and $\tau_{out} = 4.3$). (**Left column**) represent the order parameters for the whole network and each module and NMI versus frequency. The gray dashed lines are the global order parameter and NMI for the network with $\tau_{in} = 2$ and $\tau_{out} = 4.3$, (figure S1, the first row) and are plotted for the eye guide. (**Middle column**) show global and modules' order parameters and the phase differences between modules versus time. Three snapshots of the phase of the oscillators in the unit circle have been also shown. (**Right column**) indicate the connectivity matrices and color codes used to identify the values of time delays.

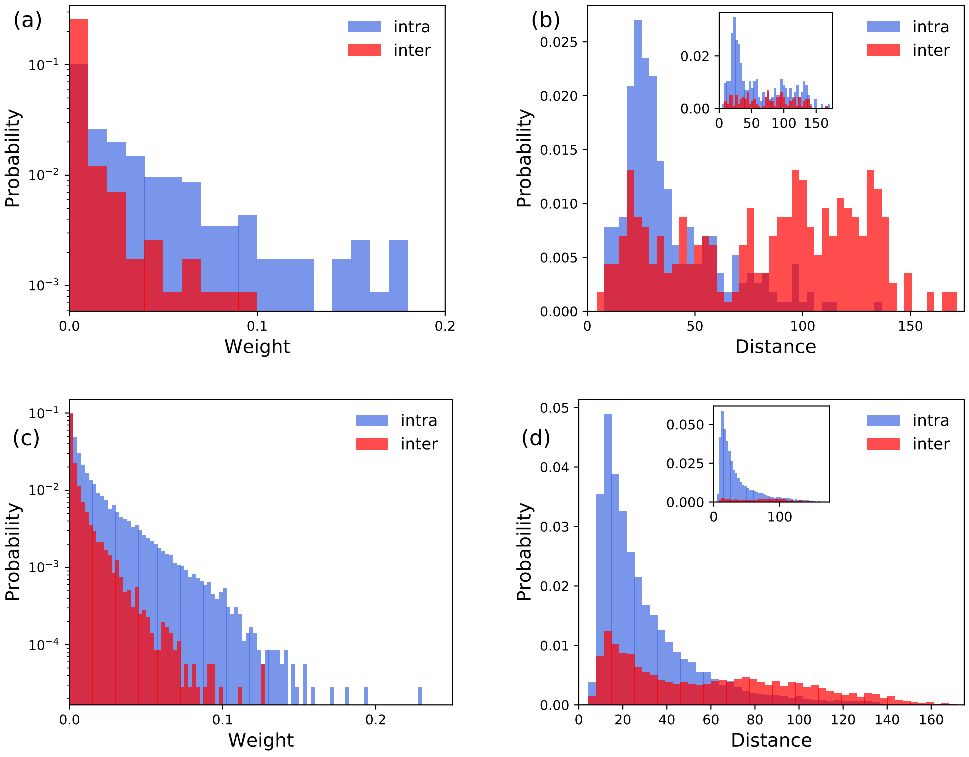


Figure S5: The distribution of distance of nodes and connection strength in the human connectomes (Hagmann et al., 2008) with **(a, b)** 66 and with **(c, d)** 998 nodes. The inset panels in **b** and **d** show the distribution of distance in level 2 of hierarchy (two hemispheres).

### 2 Hierarchical modular structure of the human connectome

Figure S6 represents human connectome in two different versions. A high-resolution matrix with 998 nodes and network (Hagmann et al., 2008) with 66 nodes that is downsampled from the previous one (Cabral et al., 2011). In this paper, we used both versions in our calculations.

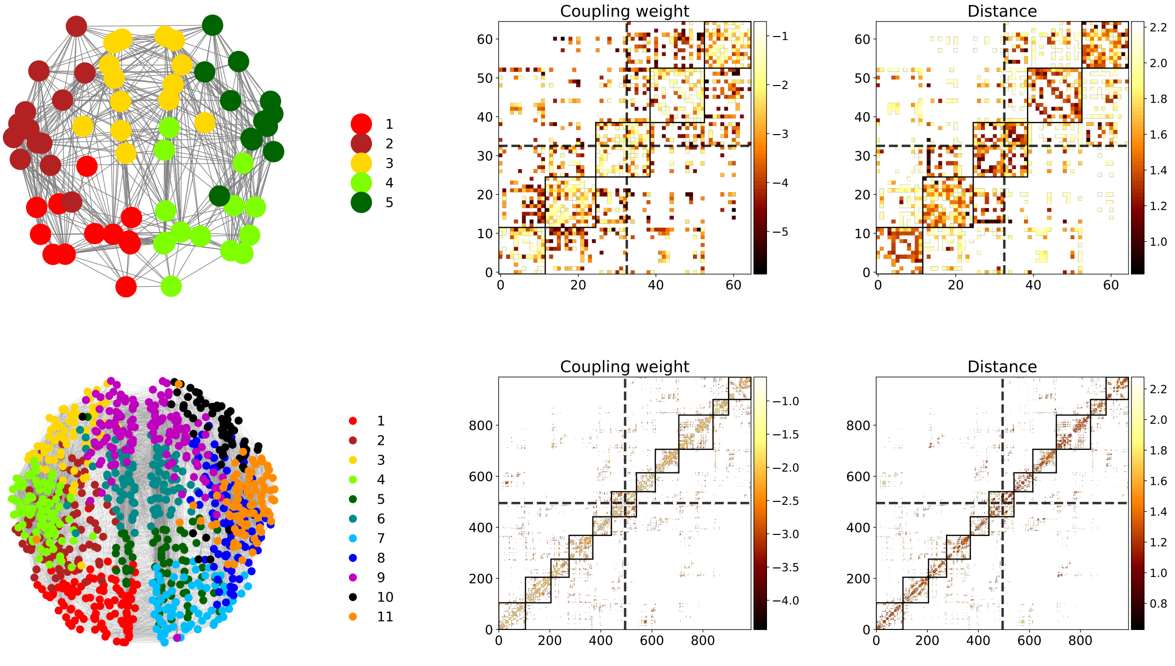


Figure S6: The representation of the human connectome with 66 (Upper row) and 998 (Lower row) nodes (Hagmann et al., 2008). In the graph representation of the brain networks the nodes in different modules are distinguished by colors. The connectivity matrices have been shown according to the coupling weights and distances. The color-bar scale is $log_{10}$. The modules are presented by black rectangles and ordered by their numbers from top to bottom, keeping the partition into the 2 hemispheres (dashed lines)

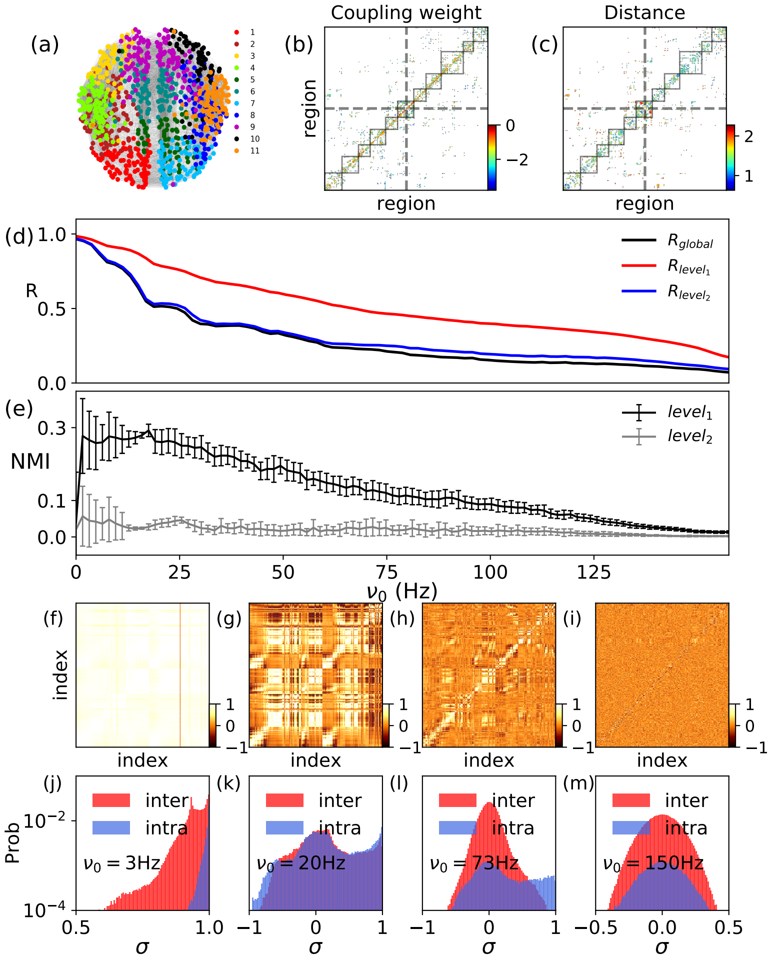

Figure S7: **The human cortex with 998 nodes.**(**a**) The graph representation of the human cortex. The colors distinguish 11 communities. (**b**) The connection strength matrix, the color-bar has $\log_{10}$ scale. The weights were normalized so that $0 \leq W_{ij} \leq 1$. (**c**) The distance between regions given as the average length of the fibers connecting pair of regions with $\log_{10}$ scale. **(d)** The time average of global and local order parameters of clusters in level 1 and 2 versus $\nu_0$ at couplings $K/N = 0.5$ where N is the number of nodes. **(e)** The NMI versus $\nu_0$ for $K/N = 0.5$. **(f-i)** The correlation matrices ($C$'s) at the same coupling $K/N = 0.5$ and different frequencies $\nu_0 =$ (3, 20, 73, 150) Hz. **(j-m)** The distribution of correlation values corresponding to panel (f-i), respectively. The results were obtained by averaging over 40 different sets of initial phases.

Table S1: Properties of each cluster in the human connectome with 998 nodes (989 after removing isolated nodes) and 17865 edges. From this number 11926 edges are inside clusters and 5939 edges between them. Size shows the number of nodes, $e_{in}(e_{out})$ is the number of edges in (between) the clusters, the next one is the density of edges and $w_{in}(w_{out})$ is the normalized sum of edge weights in (between) the clusters. $< \tau_{in} >$ and std($\tau_{in}$) are the mean value of delay and standard deviation of delays, respectively.

| index | size | density | $e_{in}$ | $e_{out}$ | $w_{in}$ | $w_{out}$ | $\tau_{in}$[ms] | std($\tau_{in}$)[ms] |
|---|---|---|---|---|---|---|---|---|
| 1 | 105 | 0.19 | 1048 | 1067 | 0.06 | 0.01 | 4.8 | 3.1 |
| 2 | 100 | 0.27 | 1345 | 989 | 0.07 | 0.03 | 6.8 | 4.7 |
| 3 | 71 | 0.30 | 757 | 1039 | 0.08 | 0.03 | 5.5 | 3.5 |
| 4 | 93 | 0.27 | 1161 | 1004 | 0.06 | 0.02 | 5.4 | 3.2 |
| 5 | 73 | 0.27 | 705 | 1380 | 0.07 | 0.02 | 6.9 | 5.3 |
| 6 | 98 | 0.33 | 1553 | 1543 | 0.07 | 0.03 | 8.7 | 7.0 |
| 7 | 74 | 0.26 | 702 | 842 | 0.07 | 0.02 | 4.3 | 2.5 |
| 8 | 92 | 0.28 | 1170 | 1003 | 0.07 | 0.03 | 7.0 | 4.7 |
| 9 | 134 | 0.21 | 1855 | 1388 | 0.06 | 0.03 | 5.7 | 5.1 |
| 10 | 61 | 0.34 | 629 | 820 | 0.08 | 0.02 | 4.7 | 3.5 |
| 11 | 88 | 0.26 | 1001 | 803 | 0.06 | 0.02 | 5.3 | 3.9 |
